# Metabolic phenotyping of marine heterotrophs on refactored media reveals diverse metabolic adaptations and lifestyle strategies

**DOI:** 10.1101/2022.01.07.475430

**Authors:** Elena Forchielli, Daniel Sher, Daniel Segrè

## Abstract

Microbial communities, through their metabolism, drive carbon cycling in marine environments. These complex communities are composed of many different microorganisms including heterotrophic bacteria, each with its own nutritional needs and metabolic capabilities. Yet, models of ecosystem processes typically treat heterotrophic bacteria as a “black box”, which does not resolve metabolic heterogeneity nor address ecologically important processes such as the successive modification of different types of organic matter. Here we directly address the heterogeneity of metabolism by characterizing the carbon source utilization preferences of 63 heterotrophic bacteria representative of several major marine clades. By systematically growing these bacteria on 10 media containing specific subsets of carbon sources found in marine biomass, we obtained a phenotypic fingerprint that we used to explore the relationship between metabolic preferences and phylogenetic or genomic features. At the class level, these bacteria display broadly conserved patterns of preference for different carbon sources. Despite these broad taxonomic trends, growth profiles correlate poorly with phylogenetic distance or genome-wide gene content. However, metabolic preferences are strongly predicted by a handful of key enzymes that preferentially belong to a few enriched metabolic pathways, such as those involved in glyoxylate metabolism and biofilm formation. We find that enriched pathways point to enzymes directly involved in the metabolism of the corresponding carbon source and suggest potential associations between metabolic preferences and other ecologically-relevant traits. The availability of systematic phenotypes across multiple synthetic media constitutes a valuable resource for future quantitative modeling efforts and systematic studies of inter-species interactions.

**Importance:** Half of the Earth’s annual primary production is carried out by phytoplankton in the surface ocean. However, this metabolic activity is heavily impacted by heterotrophic bacteria, which dominate the transformation of organic matter released from phytoplankton. Here, we characterize the diversity of metabolic preferences across many representative heterotrophs by systematically growing them on different fractions of dissolved organic carbon (DOC). Our analysis suggests that different clades of bacteria have substantially distinct preferences for specific carbon sources, in a way that cannot be simply mapped onto phylogeny. These preferences are associated with the presence of specific genes and pathways, reflecting an association between metabolic capabilities and ecological lifestyles. In addition to helping understand the importance of heterotrophs under different conditions, the phenotypic fingerprint we obtained can help build higher resolution quantitative models of global microbial activity and biogeochemical cycles in the oceans.

## Introduction

Three quarters of Earth’s surface is covered in water, making the ocean the biggest continuous environment and home to extraordinary biodiversity (1). In stark contrast to terrestrial biomes, approximately seventy percent of the biomass in marine ecosystems is microbial (vs 96% plant on land) (2); accordingly, their sub-microscale processes have global-scale consequences on ecosystem services critical to human society (3). Half of the Earth’s annual primary production is accomplished in the surface ocean by phytoplankton that harvest light to fix carbon dioxide (3–5). Heterotrophic bacteria are important players in these processes, as they impact carbon cycling in the marine environment along at least two main axes: first, phytoplankton function is modulated by interactions with heterotrophs in ways that drastically affect primary productivity (6–9). For example, heterotrophic bacteria provide nutrients essential for the long-term survival of some phytoplankton (10–13). Secondly, heterotrophs dominate the transformation of organic matter released from phytoplankton, and their metabolic activity ultimately determines the fate of organic carbon in the marine environment (6, 14).

The marine dissolved organic carbon (DOC) pool is the primary source of organic carbon for marine heterotrophs, containing an estimated 0.2 Pg of labile organic compounds, which can be readily metabolized by these bacteria (15). Analyses of bulk seawater reveal the enormous complexity and heterogeneity of marine DOC, enumerating a minimum of tens of thousands of distinct organic compounds (16–18). As the primary suppliers of marine DOC, phytoplankton transfer approximately 50% of their photosynthate to heterotrophs (4, 19) via a variety of active and passive mechanisms, including leakage (20), exudation (21), photosynthetic overflow (22), and cell death caused by viral lysis and protist grazing (23, 24). A number of studies indicate that the taxonomic composition of microbial communities is influenced by the provenance of organic material, suggesting that individual heterotrophs vary in their ability to utilize broadly-defined classes of macromolecules (25, 26) and that these metabolic preferences may contribute to community structure (27–31). For example, heterotrophs associated with phytoplankton have shown preferences for amino acids, small sulfur-containing compounds, and one-carbon compounds (32–36). However, little is known about the specific preferences of individual heterotrophs for individual classes of compounds, making it difficult to understand the role that specific clades play in utilizing DOC in different environments. This limited knowledge also makes it challenging to lay the groundwork for mechanistic models that could help explain how such interactions shape the phylogenetic and functional composition of the community. Identifying the metabolic links between DOC and heterotrophs is key to understanding the ecological drivers underpinning carbon cycling in ocean ecosystems. This may also help improve global-scale models of marine microbial processes, where heterotrophs are often assumed to perform overall similar metabolic tasks (37, 38), an assumption likely reasonable for certain goals (e.g. modeling global primary productivity) but not others (e.g. modeling community composition or genetic capacity).

In principle, information on the substrate preferences of different heterotrophs could be inferred from their genomes; however, sequencing data alone has demonstrated a limited ability to predict microbial phenotypes and community functions in practice (39, 40). For example, in a recent survey of human gut bacteria, metabolic models recapitulated growth for only 10 of the 40 strains tested, suggesting that genomic information combined with knowledge from the literature are insufficient to describe bacterial metabolic complexity (41). Alternatively, by measuring the growth properties of individual strains on specific carbon sources, one can infer phenotypic profiles that provide direct insight into metabolic preferences and growth strategies. These types of measurements are increasingly performed to characterize microbial collections from different biomes (42, 43). Ideally, one would want to analyze phenotypic profiles in conjunctions with the organisms’ genomes in order to obtain insight into the genes and pathways that confer these preferences. Performing these types of measurements on well-defined synthetic media, in addition to enabling inferences regarding the specific metabolic capacities of microorganisms, could also help inform quantitative models (44, 45).

Here we report the generation and characterization of a collection of 63 heterotrophic bacteria representative of many major marine clades, and a systematic analysis of their metabolic preferences. By designing simplified media that capture different fractions of the molecular components of marine DOC, we sought to characterize the metabolic properties of different representative heterotrophs, i.e. how they grow on different conditions that represent different axes of DOC. We next analyzed the phenotypes obtained in order to understand whether these metabolic phenotypes are well captured by phylogeny or other genome-encoded properties. We suggest that our analysis could contribute to helping reduce the complexity of the ocean microbiome to a set of computationally and experimentally tractable variables that can be interrogated with mathematical models, controlled experiments, and extended to complex natural communities to explain aspects of their behavior.

## Results and Discussion

### Growth of 63 heterotrophs on refactored media provides an atlas of their metabolic preferences

In order to generate a collection of strains representative of major marine lineages common to both the global oligotrophic and temperate oceans, we collected 63 heterotrophic isolates from different sources (Supplementary table S1). Strains were selected via a comprehensive genomic analysis conducted in a recent effort (46), in which over 400 high-quality reference genomes were clustered into functional groups using a trait-based approach focusing on metabolism and microbial interactions. Representative strains were chosen from each functional cluster, applying the additional criteria that they be culturable in standard laboratory conditions (at 26°C in Marine Broth), and have a BSL1 rating. In addition, we also included several non-marine model strains to serve as a benchmark for our experiments. Overall, the culture collection includes representatives of 5 phyla and 29 families.

We used this collection of heterotrophic bacteria to address, through single strain phenotyping, the question of whether different clades are geared towards efficient degradation or preferred utilization of specific subsets of DOC molecules. Many of the library strains had been reported to grow in undefined complex media, such as marine broth. Using the composition of marine broth as a scaffold, we refactored yeast extract and peptone (the primary carbon sources in marine broth) into eight types of organic carbon (hereafter referred to simply as “carbon classes”): peptides, amino acids, lipids, disaccharides, organic acids, neutral sugars, amino sugars, and acidic sugars (Figure 1A). For some classes, we selected specific compounds based on their reported presence in marine DOC (47–49). For example, in the amino sugar class, N-acetylglucosamine was chosen because it is a known degradation product of chitin, the primary cell wall component of many marine organisms (50). All refactored media were calibrated to contain the same total mass of carbon sources, as well as an excess of nitrogen, phosphorus, sulfur, salts, trace metals, and vitamins (Supplementary table S2). Together with a negative control lacking added organic carbon, a medium containing all carbon classes, and Marine Broth itself, we tested a total of 11 conditions using growth assays in 96 well plates (Figure 1B).

**Figure 1.**
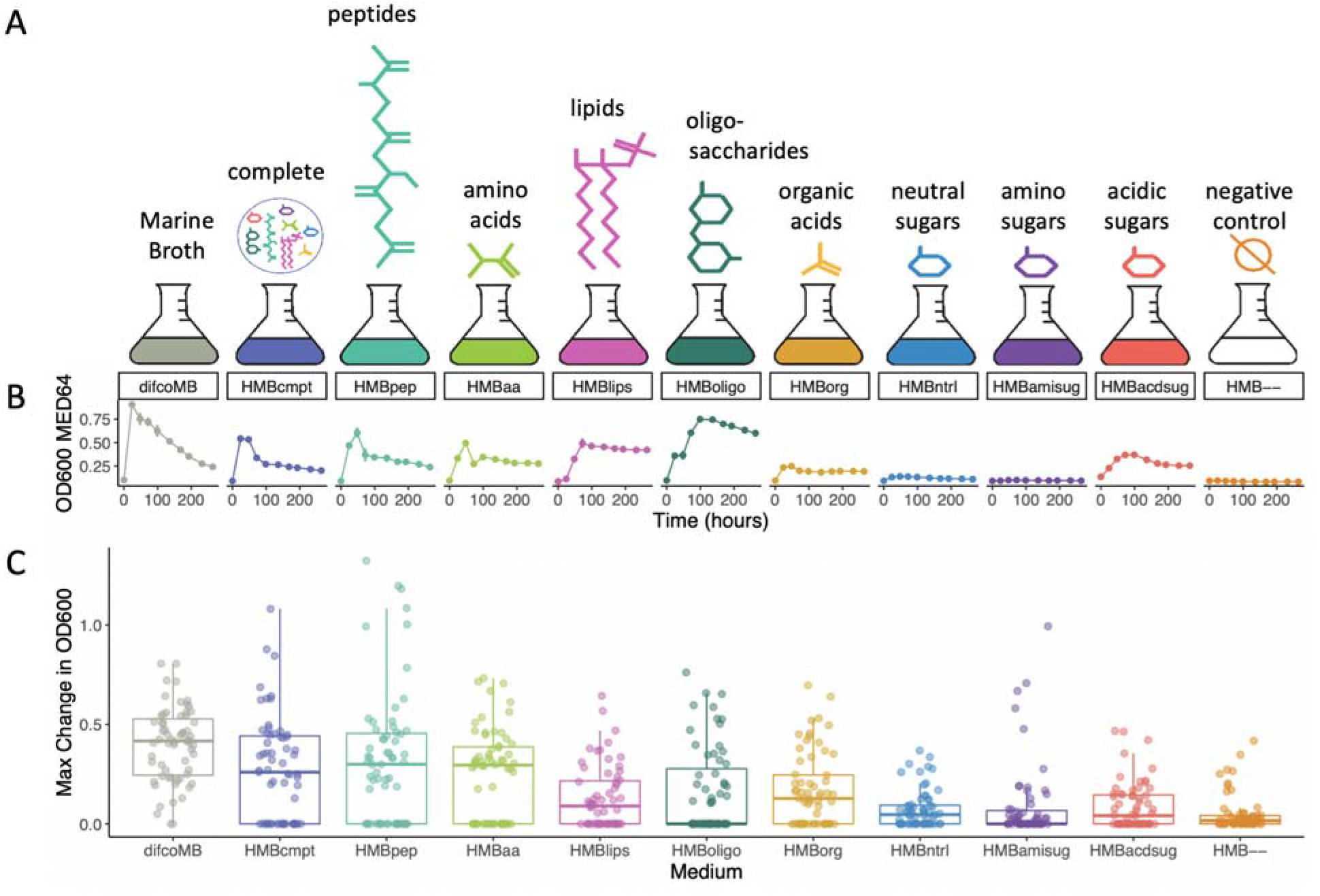
An overview of the experimental setup. Each strain in the heterotroph library was grown individually on each of the 11 different media. **(A)** Difco Marine Broth was refactored into 8 carbon classes, which were used to make defined media. The set of media used in the experiment also includes a medium containing all carbon categories (“complete”) and a negative control lacking added carbon. **(B)** Growth was assayed by measuring the OD600 over the course of 264 hours; *Alteromonas mediterranea* MED64 growth curves are shown here as an example. **(C)** MaxOD, defined as the maximum OD600 reached during the growth process (relative to the culture at time zero) was determined for each strain under every possible medium. Here, for each medium (x-axis) we visualize the MaxOD of each organism as a dot (with small random displacement on the x-axis for ease of visualization). The box represents the median and quartiles of the MaxOD values. Note that some strains reproducibly displayed non-negligible growth on the medium with no carbon added (see Table S7 for p-values). The high reproducibility between experiments (Figure S1) likely rules out contamination. Carry-over of nutrients from the starter culture is also unlikely to explain this, since these strains grew to a lesser magnitude on other media (unless inhibition by carbon in those other media is a contributing factor). We cannot exclude the possibility that these strains are capable of utilizing compounds other than the added carbon sources (e.g. vitamins) or internal carbon stores for growth, or that they possess autotrophic capabilities.

All of the 63 marine heterotroph strains grew reproducibly on at least one of the defined media, and OD measurements were almost identical across technical replicates and highly consistent across biological replicates (Figure 2, Supplementary Figure S1). For subsequent analyses we focused on the maximal OD (max OD) observed along the curve of each strain relative to time zero, which we took as a proxy for the efficiency with which each organism can produce biomass on a given carbon category (referred to henceforth also as “productivity”). Note that another important metric, the growth rate at log phase is strongly correlated with maxOD (adjusted R-squared = 0.71, p-value < 2.2×10^-16^, Supplementary Figure S2). Despite containing the same mass of carbon source and supporting roughly similar numbers of strains (Supplementary Figure S3), the different media lead to very different degrees of productivity. Given that neutral sugars, such as glucose and arabinose are classically used as preferred carbon sources in bacterial growth experiments, we were surprised to observe that media having peptides and amino acids as the main carbon sources supported the highest amount of biomass growth, while the neutral sugar medium was among the lowest supporters of biomass change (Figure 1C).

**Figure 2.**
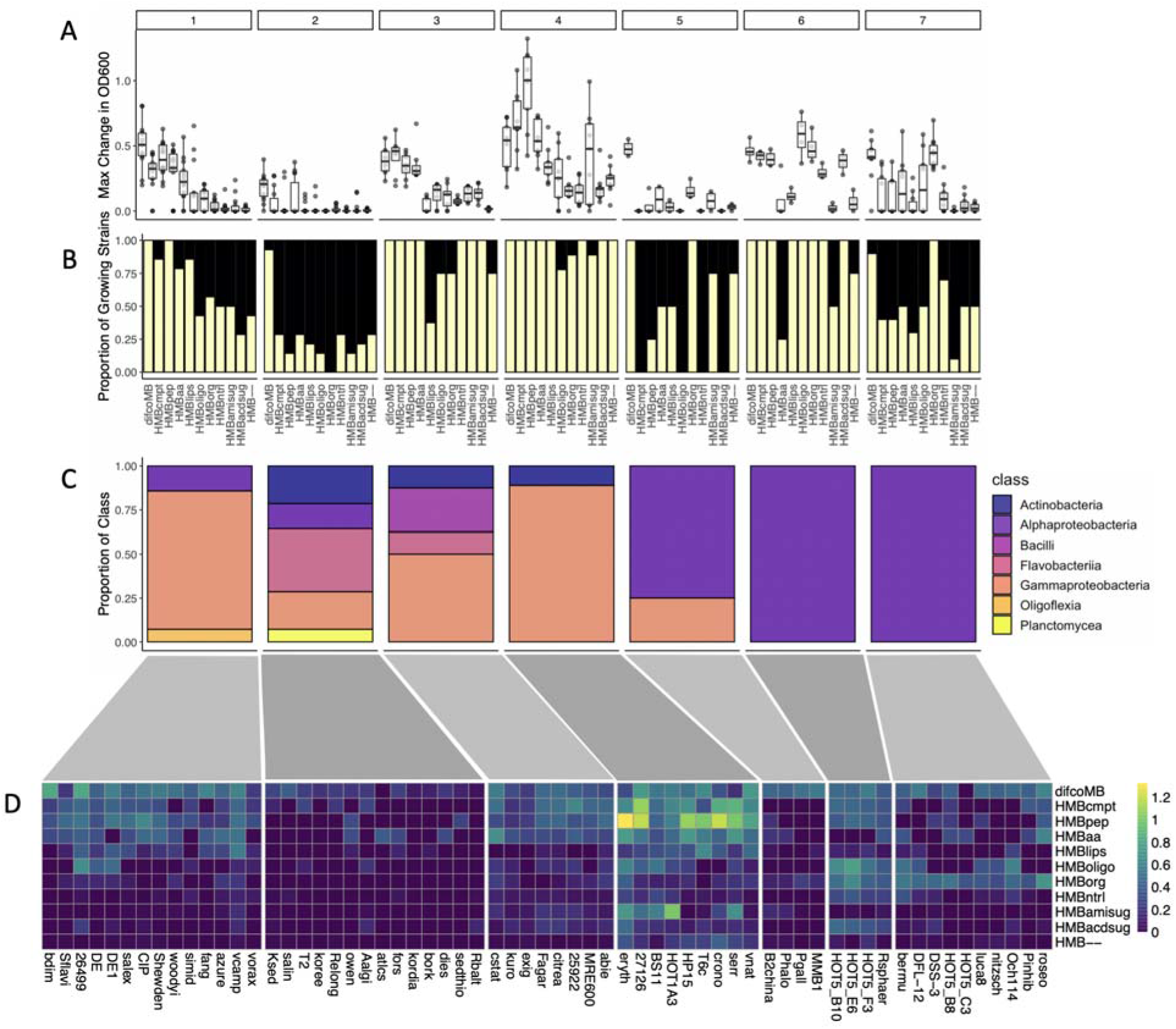
Clustering of growth profiles (defined as the vector of MaxOD values for each organism across all media) reveals metabolic signatures of the heterotroph library. **(A)** A representation of MaxOD for each cluster. Each subplot (numbered from 1 to 8) shows a boxplot of the MaxOD across media (similar to the one of Fig1C), but restricted to the organisms belonging to the corresponding cluster. For example, cluster 4 includes organisms which grow very well on the peptide-containing medium (HMBpep). **(B)** The proportion of strains (fraction, out of the total 63 strains) determined to have positive growth (see Methods) on each medium, for each of the clusters. For example, in cluster 4, all species grow on the first four media. **(C)** The proportion of taxonomic groups represented in each cluster (at the class level, see color legend to the right on the panel). **(D)** Clustered heatmap depicting the MaxOD for each strain/medium combination. For each cluster, the heatmap (media, y-axis by organism, x-axis) shows the MaxOD value (see color code to the right of the panel).

One aspect of the above results that is worth reflecting on, is the significance of this pattern in the context of prior observations. It has been suggested in the literature that marine heterotrophic bacteria achieve greater biomass yield and growth rates on amino acids compared to media containing mono or polysaccharides as the sole carbon source (51, 52). The reason for this is unclear, but field studies suggest that nitrogen limitation relieved by the presence of additional nitrogen in amino acids does not explain this discrepancy (52). We did not observe increased growth on the amino sugar-containing medium (among the lowest growing), which contains considerably more nitrogen than the non-amino media. Furthermore, a regression analysis showed that there is only a weak relationship between the change in OD and amount of nitrogen in the medium (adjusted R-squared = 0.069, p-value = 1.25×10^-9^, Figure S4). Thus, nitrogen abundance is likely not the main or only reason for the extensive bacterial growth on amino acids, and other explanations could involve the energetic advantage of using pre-formed amino acids compared to their biosynthesis (53). Note also that some organisms display small but reproducible non-zero growth on the negative control medium, possibly due to the ability to utilize other media components or internal carbon storage molecules for growth (see also Figure 1).

### Organisms cluster into two main groups and finer structures based on metabolic preferences

We captured each strain’s metabolic phenotype by compiling a growth profile, which consisted of a vector of the maximum change in OD achieved by the strain on each medium. Using a Gaussian mixture model (see Methods), these 63 growth profiles could be divided into seven clearly distinct clusters (Figure 2) The clusters can be described in terms of unique metabolic “signatures”, i.e. distinct sets of carbon classes on which the strains achieved similar biomass growth.

At a very broad level, six clusters seem to partition into two categories of metabolic preferences: one group (clusters 1, 3, and 4) includes organisms that grow robustly on amino acids and relatively poorly on organic acids (Figure 2A, B); these clusters are enriched in Gammaproteobacteria (squared standardized Pearson residual for chi-square test > 4; see methods, Supplementary Table S3). In contrast, clusters 6 and 7 are comprised of organisms that produce significantly more biomass when grown on the organic acid medium compared to the amino acid medium (Figure 2A, B, Supplementary Table S4); these clusters are strongly enriched in Alphaproteobacteria (squared standardized Pearson residual for chi-square test > 4; see methods, Supplementary Table S3). Cluster 5 follows a similar pattern to clusters 6 and 7, but the magnitude of growth is lower and the observed differences are not statistically significant (Figure 2A, B). The remaining cluster (2) is highly diverse phylogenetically, albeit strongly enriched for Flavobacteriia and Actinobacteria (squared standardized Pearson residual for chi-square test > 4; see methods, Supplementary Table S3). As a whole, the strains in cluster 2 do not grow robustly on any media type, which may indicate a requirement for growth factors or environmental conditions not represented in the media tested.

In addition to these broad-scale patterns, the strains in several of the clusters show more nuanced differences in their growth phenotypes. Firstly, although the strains in cluster 4 are capable of growing on organic acids, the magnitude of growth is greatly reduced compared to the media containing amino acids (Figure 2A, Supplementary Table S4). Interestingly, these strains consistently grow to a higher OD on the negative control medium compared to the organic acids, although this difference is not statistically significant (Figure 2A, Supplementary Table S4); this might suggest that organic acids actually inhibit their growth. In contrast, the strongest growth on organic acids is observed for clusters 6 and 7, which also display a limited and varied ability to grow on amino acids. Curiously, only 4 of the 18 strains in clusters 5, 6, and 7 grow appreciably on both peptides and amino acids (Figure 2B); the remaining 14 either grow on one or neither of these two media. Since growth on peptides requires the ability to take up and incorporate amino acids, it is unlikely that these strains lack this capability. Rather, it is possible that some of the amino acids negatively affect growth at the concentrations employed here (54). Overall, except for cluster 2, which seems to include strains with a common pattern of low OD but no specific preference for carbon classes, the heterotrophs’ metabolic functions are thus primed for optimized growth on amino acids or organic acids, but not both.

### Phylogenetic and metabolic distances correlate poorly with metabolic preference distances

The fact that phylogeny does not seem to completely explain the metabolic phenotype clusters can be explored in a quantitative way by plotting the phylogenetic distance as a function of distance in phenotype space. Specifically, we asked whether strains that are more closely related to each other phylogenetically are also more similar in their growth profiles across the different media we tested (see Methods). Regression analysis found an extremely weak relationship between the two variables (Figure 3A, adjusted R-squared = 0.001, p-value = .04), indicating that the differences in phylogenetic distances between strains explained an insignificant proportion of the variation in distance between growth profiles.

**Figure 3.**
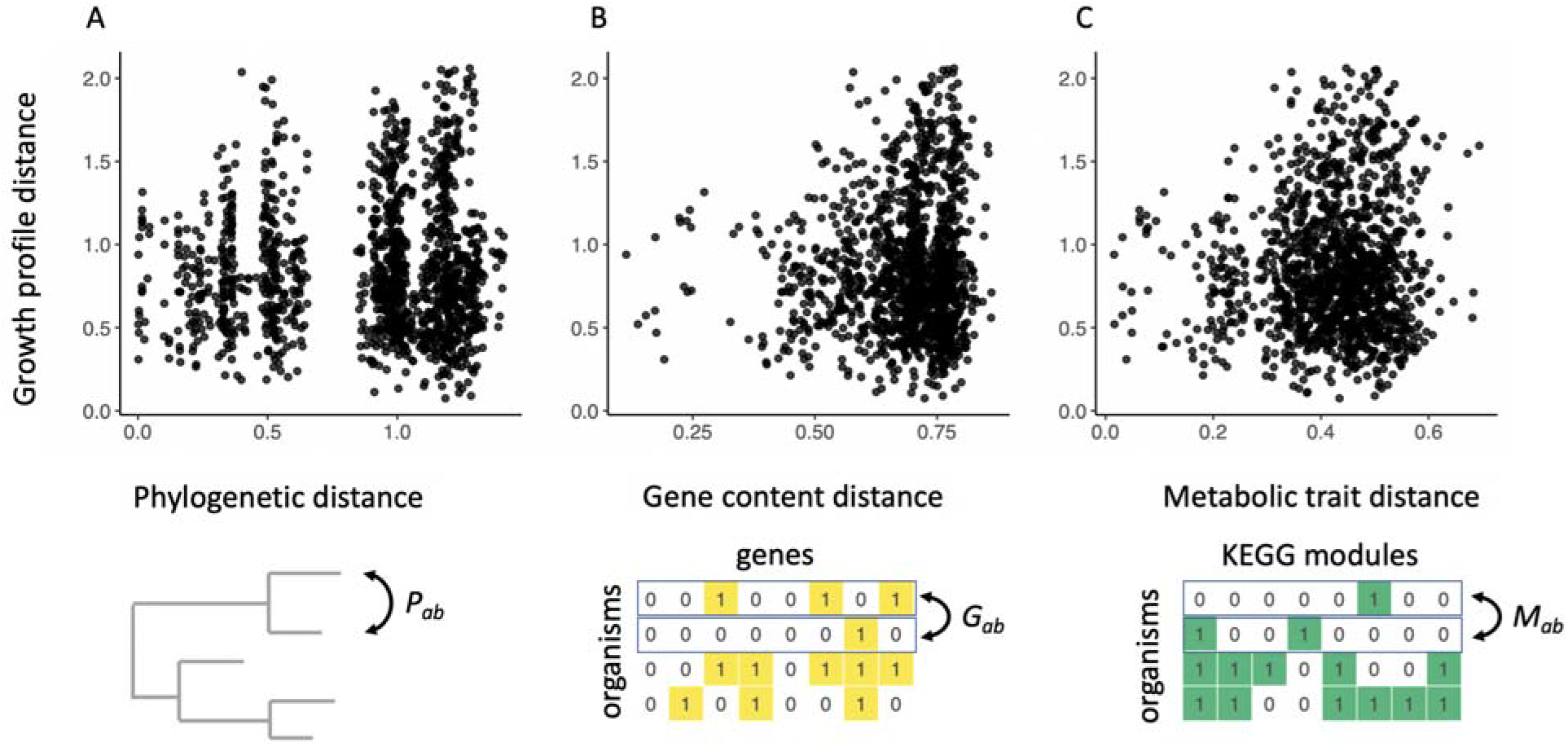
Genome-based distances between all pairs (a,b) of strains, i.e. phylogenetic distance *P_ab_* **(A),** genome-wide gene content distances *G_ab_* **(B)** and KEGG module distance *M_ab_* **(C)** (see Methods) are compared to the phenotypic distance (y-axis in all panels) between the corresponding growth profiles shown in Figure 2D. All of the genome-based distances are but poor predictors of the distance between the corresponding growth profiles.

Subsequently, we asked whether a metric other than phylogenetic distance, e.g. the difference in gene content, would correlate more strongly with phenotypic distance. Specifically we compared distances between growth profiles to distances between genomes represented as binary sequences of genes (see Methods). Linear regression revealed an inconsequential association between growth profile distance and genome distance (Figure 3B; adjusted R-squared = 0.004, p-value = .0005). The same was true when we compared the growth profile distance to the distance between KEGG modules, which better approximates the metabolic distance between strains (Figure 3C, adjusted R-squared = 0.004, p-value = .0004) (46). Taken together, global genomic data, at least with the metrics used so far, do not seem to capture the differences in metabolic phenotypes observed between strains. One possible explanation for our observations is that the genes key in differentiating growth on various carbon sources are “drowned out” by the noise of uninformative genes contained in the genomes; it’s not clear that the genes that matter rise above genes that have no effect in this analysis.

### Genes and pathways associated with growth on specific media reflect different metabolic and ecological strategies

Given the poor correlation between metabolic distance and gene content or phylogenetic distance, we asked whether a more informative relationship could be revealed by examining whether the presence of individual genes or sets of genes (pathways) is associated with growth on each medium. For each medium, we calculated the correlation (r) between growth and each gene in the library pangenome (see Methods, Figure 4A). Looking at the list of individual significantly correlated genes (supplementary table S5), one clear pattern that emerges is that 386 of the 1076 most strongly correlated genes (absolute value of r > 0.5) are associated with growth on the organic acids medium. Conversely no significant individual genes emerged for growth on the amino acid medium, despite the fact that this medium supports high growth in a number of strains. One possible interpretation of this difference is that while organisms growing on organic acids tend to use a narrow and fairly coherent set of pathways, the organisms that grow on the amino acid medium are utilizing different subsets of the 20 amino acids, and thus different biological pathways (see supplementary table S2 for details). A concise view of all gene correlation scores can be visualized using PCA: as shown in Figure 4B, the organic acid medium stands out as the main contributor to the primary PCA axis, and is thus the most unique in terms of enriched genes compared to the other media. It is also interesting that the media containing additional nitrogen in the form of amino groups appear to cluster together, suggesting that genes related to the utilization of organic nitrogen partially drive this clustering.

**Figure 4.**
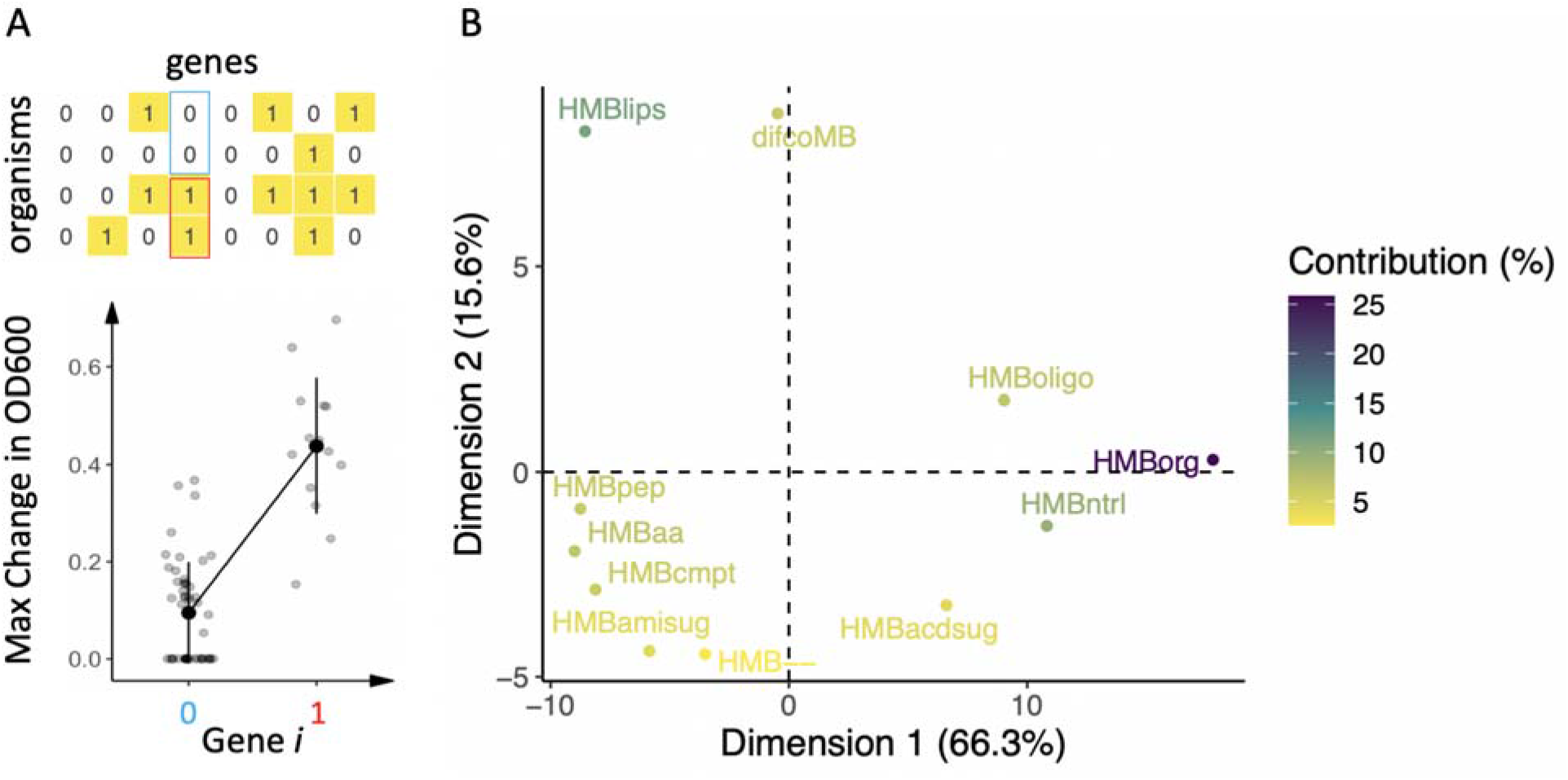
**(A)** A schematic representation of the process for generating gene-specific correlations between presence/absence of each gene and growth on a given medium. For each medium and each gene, we plot MaxOD vs. gene presence/absence for all organisms, and compute the point biserial correlation (see Methods). This gives rise to a matrix of correlations indicating how much the presence of each gene is predictive of growth (across all organisms) on a given condition. This matrix can also be viewed as a collection of row vectors (of length equal to the number of genes), each representing a condition. **(B)** Through dimensionality reduction of these row vectors (with PCA) we visualize how similar the media are to each other, in terms of the genes correlated with growth. The first two PCA axes account for 81.9% of the variance. Data points are colored according to the contribution of the samples (media) to the principal components.

In order to better identify genomic signatures associated with growth phenotypes beyond individual genes, we implemented a gene set enrichment analysis (GSEA) to identify overrepresented pathways. We mapped the set of highly correlated genes to the KEGG database (see Methods), which resulted in a ranked list of pathways for each medium (supplementary table S6); we will refer to these pathways simply as condition-specific “enriched pathways”. We next asked whether the enriched pathways are indicative of the medium under which the enrichment was identified. In other words, are the growth media enriched for pathways that metabolize the class of carbon substrate they contain?

In several cases, growth on the various media was associated with the KEGG pathways describing the metabolism of the specific compounds they contained. For example, the sugar-based media (HMBoligo, HMBntrl, HMBacdsug) are depleted of pathways involved in the metabolism of amino acids and enriched for pathways involved in galactose, starch and sucrose metabolism, and interconversions between the pentose monosaccharides and glucuronate, a degradation product of alginate (Figure 5A) (49). The media lacking sugars are not enriched for these sugar degradation pathways; instead, they share an association with the glyoxylate and dicarboxylate metabolism pathway, which can replenish sugars from amino acid precursors. In other cases, growth on a specific category of carbons is associated with pathways that can be involved in the utilization of those compounds, but deviate from the most basic expectation. For example, organisms growing on organic acids are enriched for specific portions of the Ethylmalonyl-CoA Pathway (EMC) (Figure 5A, Supplementary Figure S5A), a well-described method for the assimilation of two-carbon compounds and biosynthesis of carbohydrates from fatty acids (55). Notably, the EMC pathway is an alternative to the glyoxylate shunt (56, 57), which is used to feed anapleurotic reactions of the TCA during growth on C2 substrates, such as acetate (Figure 5B). A key enzyme in glyoxylate pathway, isocitrate lyase, is known to be absent in certain marine bacteria, such as *R. sphaeroides*, which nonetheless possess the ability to grow on acetate as a sole carbon source (58).

**Figure 5.**
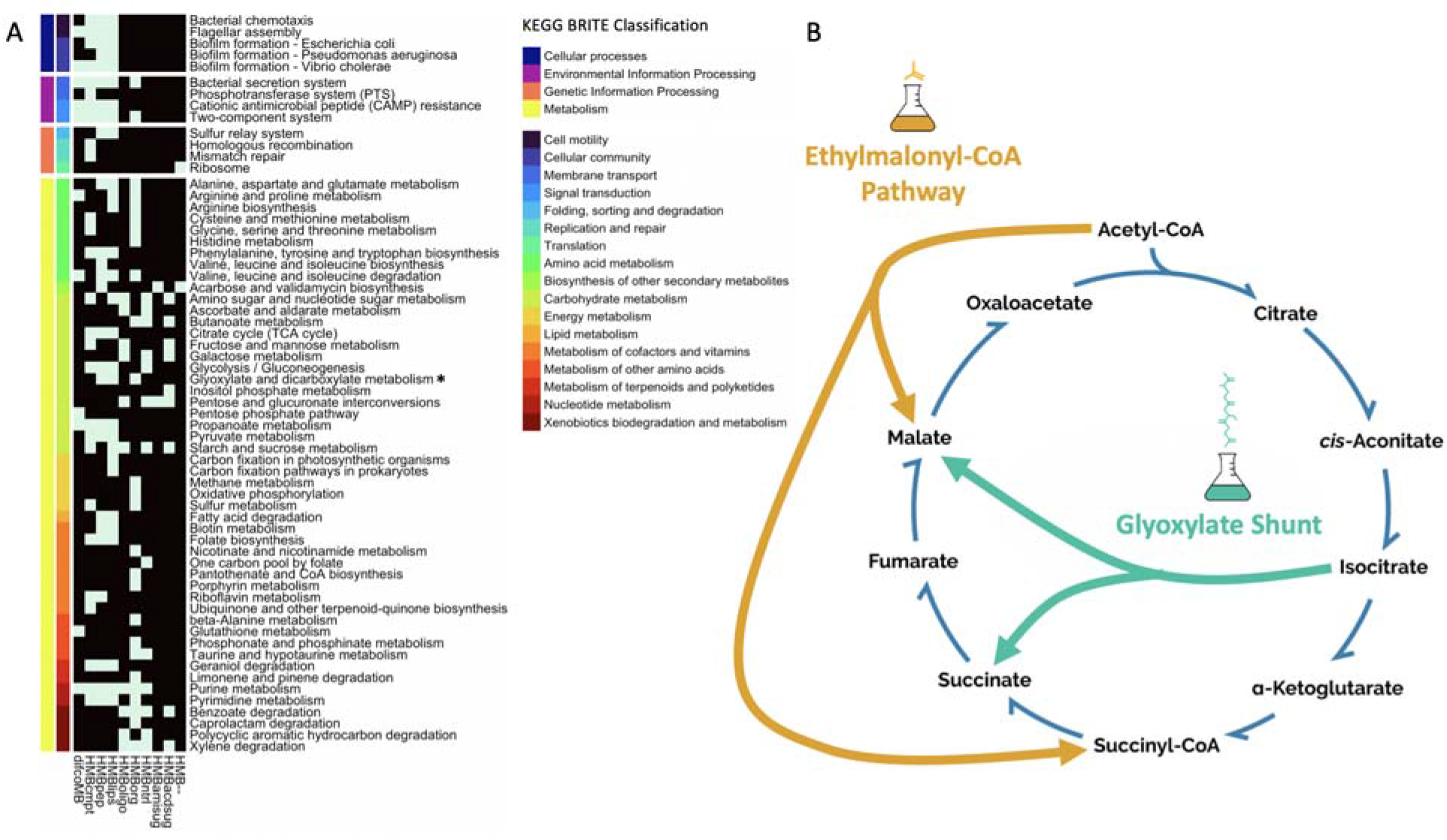
Growth on each medium is associated with specific biological pathways, as determined by a gene set enrichment analysis (see Methods). **(A)** The set of highly-correlated genes for each medium was mapped to the KEGG pathway database; for each medium, pathways that are significantly overrepresented in the highly-correlated genes are colored light blue (black indicates a lack of statistical enrichment). KEGG pathways are annotated with colors corresponding to their BRITE database classification. **(B)** One of the interesting pathways that emerges from this analysis is the Ethylmalonyl CoA pathway (EMC) pathway, which appears in KEGG as part of the Glyoxylate and Dicarboxylate Metabolism pathway (asterisk in panel A). The EMC pathway is strongly enriched for growth on organic acids. This pathway is a multifunctional pathway, known to be usable for organic acid assimilation as an alternative to the glyoxylate shunt.

Not all pathways found to be enriched can be easily associated with degradation of the carbon sources included in the medium: these pathways seem to have no direct relationship to the molecular processes associated with the utilization of the corresponding media components. For example, growth on organic acids is also correlated with the KEGG pathway for porphyrin metabolism, specifically the complete pathways for vitamin B12/cobalamin biosynthesis as well as bacteriochlorophyll a and b (Supplementary Figure S5B). On the other hand, the folate biosynthesis pathway is enriched for growth on peptides, lipids, and the complete medium (Figure 5A). In addition, growth on peptides (and lipids) is associated with pathways for motility, including chemotaxis and flagellar assembly, as well as for biofilm formation (Figure 5A).

## Conclusions

By defining a suite of carbon compound classes that together make up biomass and asking which marine bacteria grow on each class, we generated a map of the broad-scale metabolic preferences of these bacteria. This map is important because heterotrophs process much of the dissolved oceanic carbon, in ways that may depend on the emergent metabolic properties of microbial communities to which they belong. Obtaining a clear picture of the carbon utilization capabilities of individual strains in relationship to their taxonomic signature seems, therefore, an essential step towards understanding how heterotrophic bacteria impact carbon cycling in the marine environment. Understanding individual strain traits will further allow researchers to know if it is feasible and reasonable to use a few representative taxa and metabolic processes as a general effective description of heterotrophic metabolism in the oceans, e.g. for the purpose of implementing mechanistic models of communities and global ocean biogeochemical cycles. At the same time, irrespective of the details of the phylogeny-metabolism relationship, one can view the broad-scale approach presented here as a valuable effort towards building more informed dynamical models of ecological processes. In particular, we suggest that partitioning heterotrophs into clusters based on their preferred utilization of combinations of DOC fractions may add to existing observations of successions, and inform ecological models at desired levels of resolution. This work might help identify whether the broad-scale metabolic preferences suggested to explain microbial succession patterns in the marine environment (30) agree with the growth preferences of heterotrophic bacteria, and perhaps predict succession patterns based on heterotrophic growth. Our results suggest that these preferences are partially predictable based on phylogeny, but that extensive intra-clade variability requires alternative approaches to group bacteria based on their metabolic role in the oceans. For example, bacteria belonging to cluster 4 (including some Alteromonas and Marinoacter strains) would be highly competitive across a wide range of organic matter, as they grow fairly well on amino acids in addition to lipids, disaccharides, and (to some extent) amino sugars. In contrast, those belonging to cluster 1 seem to be more specifically tuned to amino acids.

While some of the associations between pathways and carbon class can be easily attributed to a direct enrichment of the corresponding metabolic utilization processes, other enrichment patterns point to processes (including non-metabolic ones) for which the biological connection is not obvious. We hypothesize that these “cryptic” associations between carbon classes and enriched KEGG pathways carry information about environmental adaptations, suggesting a connection between the metabolic preferences of marine heterotrophs and ecologically-relevant roles they play within a microbial community. For example, the fact that biofilm formation and motility are enriched for organisms that grow well on peptides is consistent with the notion that these strains are adapted for particle-associated lifestyles involving intermittent rapid growth in areas of patchy nutrient availability. A second example is that of bacteria growing on organic acids, which we found to be strongly enriched for the ethylmalonyl-CoA pathway, possibly reflecting the importance of this pathway in the photoheterotrophic capabilities shared by many of these bacteria. These same bacteria also display strong enrichment for vitamin B12 production (a defining characteristic of interactions phytoplankton-heterotrophs interactions (59)), corroborating the previously suggested role of these bacteria as key partners of eukaryotic phytoplankton. These findings are also consistent with genomic and experimental evidence that photoheterotrophy is a widespread metabolic strategy among certain Alphaproteobacteria clades (60, 61), such as the Rodobacters from our library. Previous studies have indicated that these strains lack the genes for carbon fixation, and are therefore unable to grow autotrophically, but use light to supplement their energetic requirements (61, 62). This agrees with our observation that these strains did not grow in the medium lacking carbon sources, although it is interesting to speculate about the possibility that the ethylmalonyl-CoA pathway could play a dual role in organic acid assimilation (as supported by the enrichment mentioned above) and CO_2_ fixation, as documented in other photoheterotrophs (63). Taken together, these cryptic associations can be interpreted as correlated adaptations with a potential ecological significance, similar to recently identified linked trait clusters (46).

While the phenotypic matrix shown in Figure 2 is generally thought in terms of its columns, representing the growth profiles of the different organisms, one can also reflect on the relevance of its rows, which display the type of communities that each DOC fraction is able to support. Notably, different carbon sources seem to support very different numbers of taxa: for example, amino acids support many different species, while amino sugars only support appreciable growth in a narrow category of bacteria. The broad utilization of amino acids by many organisms may simply reflect the fact that they may be able to directly import and use amino acids as building blocks, funneling them directly into biomass. Alternatively, it is possible that distinct sets of organisms preferentially use different individual amino acids, in a way that our current setup (where all amino acids are mixed into a single carbon fraction) would not be able to dissect. One could apply to carbon fractions a categorization similar to the one used to describe microbial species as generalists or specialists. In the same way as a species that can grow on multiple nutrients is thought of as a generalist, a carbon fraction that can support multiple clades could be called “versatile”. A carbon fraction that only supports a small number of clades (similar to a specialist organism) could be thought of as non-versatile, or “exclusive”. In the future it will be interesting to study the distributions of versatility in different environments and their ecological implications, e.g. with consumer-resource models that use statistical ensembles of random matrices to parametrize ecological models (64).

This laboratory study only provides a glimpse of the myriad factors likely to influence heterotrophic metabolic activity in the ocean, as the complete picture of microbial growth encompasses much more than the quantity and structure of carbon sources. Prior work has indicated that the elemental stoichiometry of DOC has a strong influence on microbial community function (65). While our refactored media are designed so as to contain the same amount of carbon, we cannot exclude the possibility that the elemental ratios of the defined media affect growth beyond the effect of the specific carbon sources. A compelling next step would be to examine the effect of nutrient availability on heterotroph carbon source preference. The landscape of organic matter in the ocean is far more complex than could be represented in laboratory experiments, and biotic interactions greatly influence metabolic activity. However, some of the trends we observed have been reported by previous studies, supporting the ecological relevance of our dataset (30, 66). While we do not expect the exact patterns observed in our experiments to play out in nature, our study provides a framework to continue investigating the role that heterotrophic bacteria play in carbon cycling.

## Materials and Methods

### Selection and construction of strain library

A library of 63 heterotrophic isolates representing major marine lineages common to both the global oligotrophic and temperate oceans (supplementary table S1) was assembled based on the classification into Genome Functional Clusters presented in Zoccarato et. al. (46). Strains were obtained from the sources listed in supplementary table S1. Samples were streaked on marine agar, and single colonies were subsequently picked and inoculated into Marine Broth. Stocks were prepared from liquid cultures in 50% glycerol and stored at −80°C.

### Construction of refactored media

As a first step in our analysis we compared strains according to their growth phenotypes with the goal of determining to what extent heterotrophic bacteria can utilize available marine carbon sources. We therefore took a well-known marine culture medium (Difco Marine Broth)-- on which all strains were experimentally determined to grow-- and selected a core set of macromolecular compounds likely to comprise the undefined components of marine broth: yeast extract and peptone. Eight media containing different classes of carbon sources were developed, each with multiple individual components: peptides, amino acids, lipids, organic acids, disaccharides, monosaccharides, amino sugars, and acidic sugars (supplementary table S2). A medium containing all eight classes and a negative control lacking added carbon were developed as well. The refactored media were designed to have the same mass of added carbon; in addition, nitrogen, phosphorus, and sulfur sources, as well as salts and vitamins were added in excess.

### Growth assay

Optical density was used as a proxy to assess biomass production, and thus the growth of the strains. While OD is an imperfect metric for bacterial growth, its widespread and historical use combined with the ability to measure non-invasively made its selection appropriate. Using sterile technique, strains were streaked from frozen stocks (maintained at −80°C) onto marine agar plates and placed in a 26c incubator. Cultures were grown 72 hours, then single colonies were picked and inoculated in 2 mL marine broth in 5 mL falcon tubes. Liquid cultures were grown with shaking (200 rpm) at 26c and ambient light for 48 hours. Negative controls without added bacteria were used to verify the absence of contamination. 1.5 ul of starter cultures were then inoculated into each well of a 96-well plate containing 149 ul of medium, in triplicate. Only the interior wells were utilized; edge wells contained marine broth without added bacterial cultures to reduce evaporation and check for contamination. All combinations of strains and media were tested. Plates were wrapped with parafilm, and incubated at 26c and ambient light without shaking for 264 hours. Negative controls for each medium (without added bacteria) were included in addition to edge wells. Plates were removed from the incubator individually, the lids were checked for condensation, and the optical density (600 nm) was measured in a Biotek Synergy HT plate reader (software version 3.05.11) at 26°C approximately every 24 hours). Plates were shaken prior to the read at 0 h only. Results (in the form of time series of OD600 for each well) were downloaded from the plate reader software as excel files and analyzed using the R statistical programming language (67).

### Growth Profiles

The heterotrophs selected for the library displayed widely divergent growth dynamics, and we chose to focus on their potential to generate biomass regardless of growth rate. As described below, we used the growth curves to estimate, for each strain, a growth profile across media, capturing the maximal change in optical density achieved by that strain at any time point during the growth assay. The three technical replicates for each organism (*i*) and condition (*j*) were highly similar to each other (supplementary figure S1), and were averaged at each time point to produce an average growth curve OD_ij_(t). We next identified for all pairs *i,j*, the maximum value (max_t_{OD_ij_(t)}) reached by OD_ij_(t) throughout the growth time course. The normalized version of this maximum value across all strains, maxOD_ij_ = max_t_{OD_ij_(t)}/OD_ij_(t=0), constitutes a matrix whose row *i* represents the growth profile of organism *i* across all conditions. The matrix maxOD_ij_ (i=1,…,63; j=1,…,10) is used for subsequent comparative analyses of heterotroph phenotypes. Following a Kruskal-Wallis test for each medium, the non-parametric post-hoc Dunn’s test was used to determine significant growth between the test and negative control without added bacteria for each medium. The Benjamini-Hochberg correction for multiple testing was applied, and cases with p-values less than 0.05 were ascribed the unchanged maxOD_ij_ value. Conversely, maxOD_ij_ values for strain/medium pairs that did not experience positive growth were set to 0.

### Unsupervised Clustering of growth profiles

The growth profiles (maxOD_ij_) of all 63 strains were clustered using a Gaussian mixture model implemented in the R mclust package (68), a contributed R package for model-based clustering, classification, and density estimation based on finite normal mixture modeling, abundantly used to analyze biological datasets (e.g. (69–71). Model-based clustering approaches, such as gaussian mixture modeling, provide a probabilistic alternative to traditional methods, such as k-means and hierarchical clustering; the challenges of selecting the “correct” number of clusters and clustering method are resolved by statistical model selection rather than heuristic methods (72, 73). Mclust provides functions for parameter estimation via the expectation maximization algorithm for normal mixture models with a variety of covariance structures, and functions for simulation from these models. Initialization is performed using the partitions obtained from agglomerative hierarchical clustering.

By default, Mclust applies 14 models and identifies the one that best characterizes the data. Mclust achieves this by computing, for each model, the Bayesian information criterion (BIC), which has been shown to work well in model-based clustering (Dasgupta and Raftery 1998; Fraley and Raftery 1998). Specifically, BIC was used within Mclust to identify the optimal covariance parameters (in our case VEI), as well as the optimal number of clusters (in our case, 7). See supplementary figure S6 for a comparison of BIC values obtained for all models.

A chi-square test of independence was performed to examine the relationship between taxonomic class and cluster number. The relationship between these variables was significant, (p-value = .0005). The squared standardized Pearson residuals were then examined to indicate which classes of bacteria contributed significantly to the lack of fit between the observed data and null model (the taxonomic class and cluster assignment are independent); values greater than 4 were considered statistically significant at a critical alpha value of .05 (supplementary table S3).

### Relationship between growth profiles and phylogenetic/gene content distance

We constructed a phylogenetic tree for all the strains in our collection using FastTree (74): protein sequences of 206 single-copy homologous genes were aligned, concatenated, and used to infer a maximum-likelihood phylogenetic tree. Briefly, 206 single-copy homologous proteins shared by all strains were identified and clustered using BLAST(75), HMMER(76), and MCL(77). Muscle(78) was used for alignment before the final tree was generated using FastTree(74). The cophenetic distance (*P_ab_*) between all pairs of strains was calculated using the cophenetic.phylo function of the R ape package (79).

The gene content distance (*G_ab_*) and the metabolic trait distance (KEGG module distance, *M_ab_*) for all pairs of strains were calculated as described in Zoccarato et al (46), using the Jaccard distance. For each pair of strains, the euclidean distance between growth profiles was plotted against the cophenetic distance, gene content distance, and KEGG module distance. Linear regression was performed using the lm function of the R base package (67).

### Correlations between growth and individual gene presence/absence

The point biserial correlation coefficient (r) was calculated between growth on each medium and the presence (binary) of each gene in the library pangenome. A subset of genes was taken from the library pangeome, eliminating genes shared by all strains or by fewer than 3 strains. P-values for r were adjusted using the Benjamini-Hochberg procedure, then filtered for values less than 0.05. The resulting set of r were considered the values for highly-correlated genes. The matrix of correlation values, r, was subjected to principal component analysis using the princomp function of the R base package (67).

### Pathway mapping/enrichment

Enriched pathways were obtained by mapping the sets of highly-correlated genes for each medium to the KEGG database using the R cluterProfiler package (80). P-values were calculated by the hypergeometric distribution and corrected for multiple testing using the Benjamini-Hochberg procedure. Enrichment was considered statistically significant for adjusted p-values less than 0.05.

## Data Availability Statement

All raw datasets and scripts used to generate the figures presented in this manuscript are available at https://github.com/segrelab/marine_heterotrophs.

## Author Contributions

EF, DSh and DSe designed the study. EF performed the experiments, and the statistical analysis of the data. EF and DSe wrote a first version of the manuscript, and all authors refined the final version. All authors have read and approved the final version of the manuscript.

## Acknowledgments

We are grateful to Melisa Osborne for support with the acquisition of the microbial strains involved in this study, and to several labs that kindly provided some isolates (see Supplementary Table S1). We are also grateful to members of the Segrè and Sher labs for constructive feedback on the design and execution of this project, to Luca Zoccarato for assistance with the genome analysis and metabolic trait distance, and to members of the labs of Hans-Peter Grossart, and Maren Voss for helpful and fun discussions on marine microbes. This work was supported by the Human Frontiers Science Program (grant RGP0020/2016) and the National Science Foundation (NSFOCE-BSF 1635070) to DSe and DSh. EF was partially supported by the Association for the Sciences of Limnology and Oceanography (ASLO) LOREX fellowship, NSF-OISE #1831075. DSe and EF also acknowledge support from NSF grant OCE 2019589 (Center for Chemical Currencies on a Microbial Planet).

