## Supplemental Figures S1-S6 for "Metabolic phenotyping of marine heterotrophs on refactored media reveals diverse metabolic adaptations and lifestyle strategies"

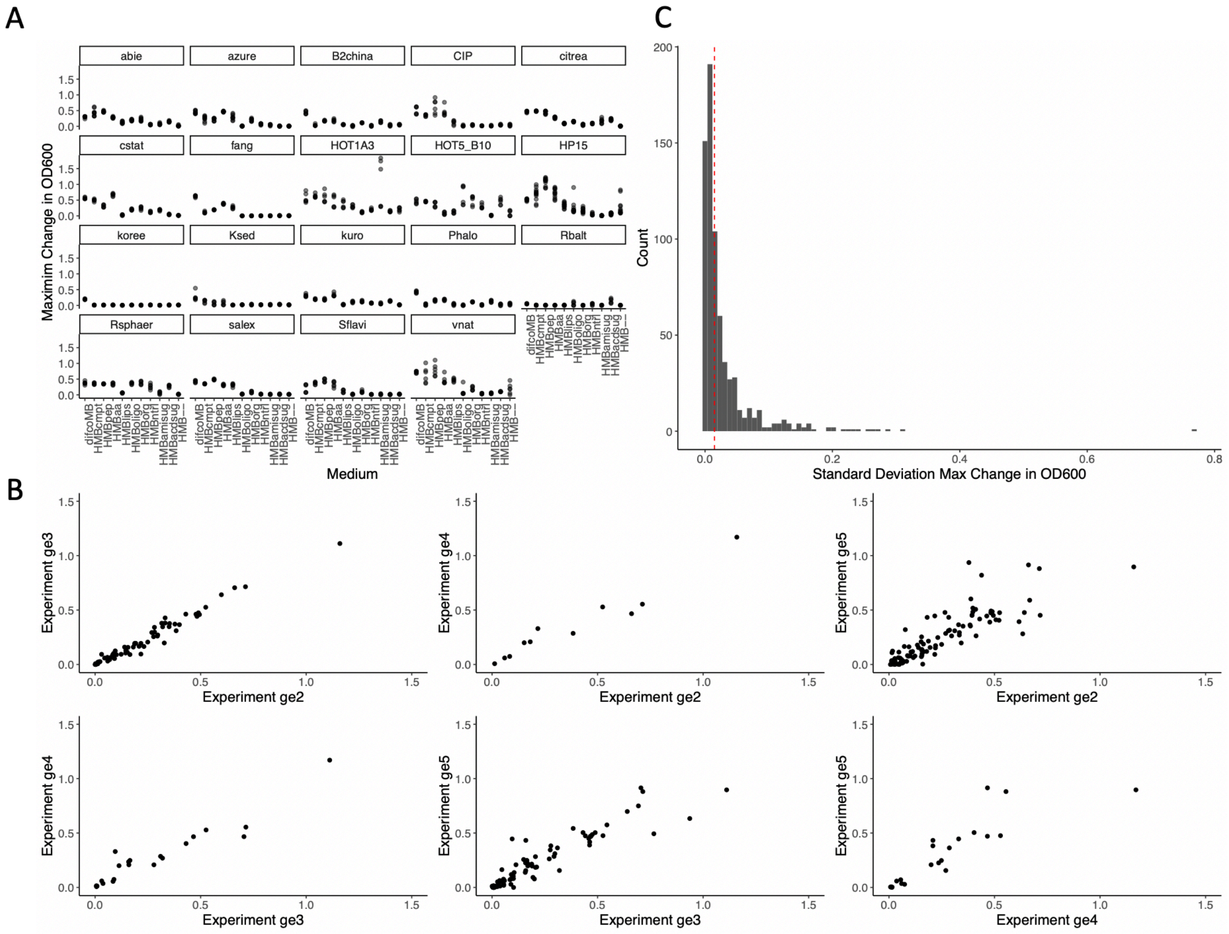


**Figure S1.** Analysis of experimental reproducibility for the growth measurements, through the comparison of biological replicates. (**A**) Each panel displays all growth measurements across the different media for a given strain. The 19 panels correspond to the 19 strains for which 4 different biological replicates (each with 3 technical replicates) were assessed. For other strains, the measurements involved 2 or 3 biological replicates (see Supplementary Data). In each panel, the box plots reflect the distributions of all the (4x3) different repeats. Spread of these repeats varies across strains and conditions, and is greater for larger OD values. (B) All MaxOD measurements for different strains on different media are compared across different pairs of biological replicate experiments. Each dot represents the average of three technical replicates. (B) The distribution of standard deviation values between technical replicates for all combinations of strains and experiments. Overall, the median of this distribution of standard deviations (~0.01, dashed red line) is much lower than the average change in OD reached across all experiments (~0.1) supporting the high reproducibility of the experimental setup.

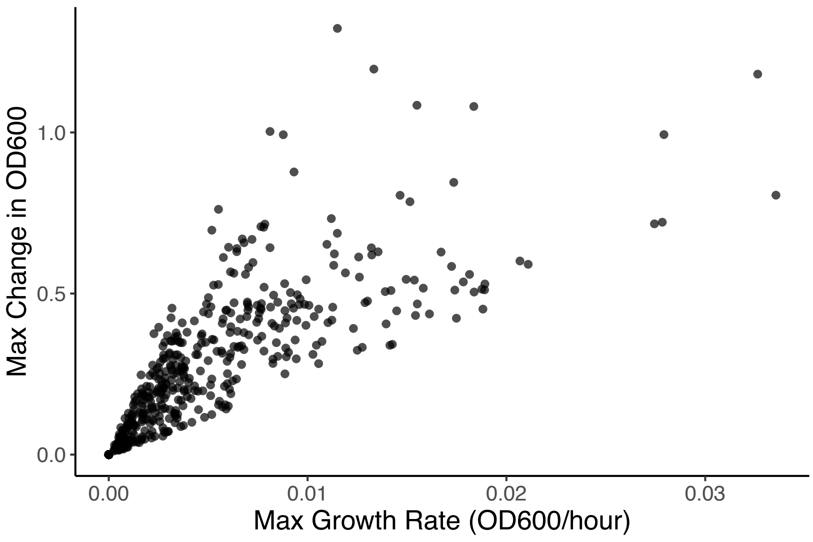


**Figure S2**. The scatter plot displays the maximum change in OD600 (MaxOD, see Methods) as a function of the maximum growth rate (OD600/hour) for all strain/medium combinations. The maximum growth rate was identified for each growth curve as the maximal slope observed across neighboring time points. There is a strong positive correlation between the two (adjusted R-squared = 0.68, p-value < $2.2x10^{-16}$).

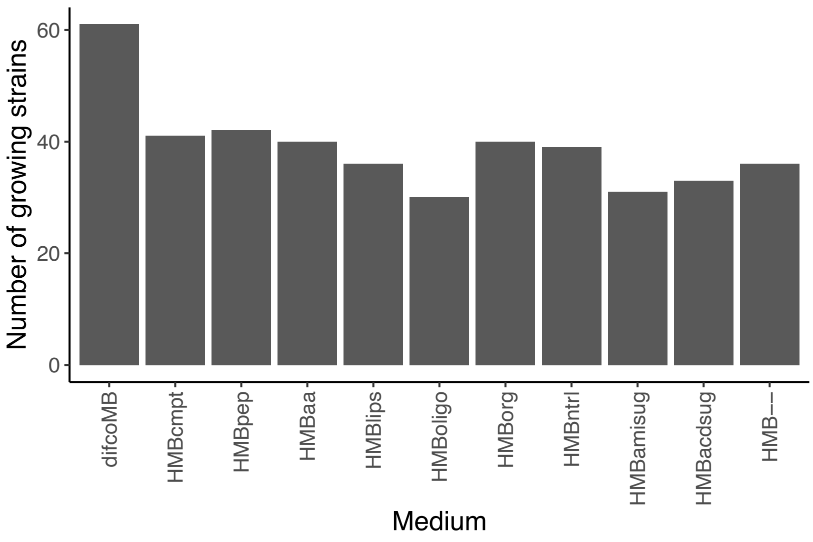


**Figure S3.** The number of strains that display positive growth on each medium. Growth was considered positive if statistically significantly greater than the negative control sample lacking added bacteria (see methods, Supplementary Table S7).

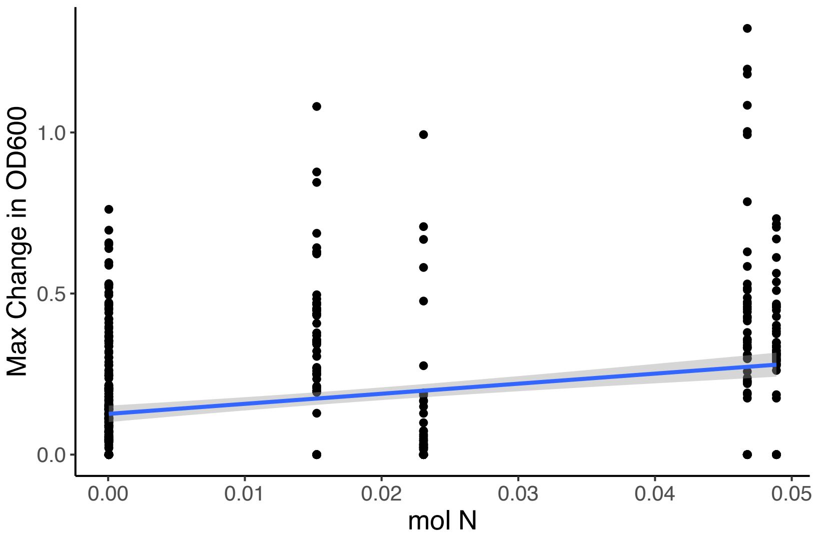


**Figure S4.** While media were built to have an equivalent total amount of carbon, they may differ in their total amount of nitrogen, potentially affecting growth patterns. Regression analysis showed however that there is a very weak relationship between growth (MaxOD) and amount of nitrogen in each medium, across all strains and conditions. Media with higher N have a statistically significant but very small increase in MaxOD (adjusted R-squared = .13, p-value < $2.2x10^{-16}$), and cannot account for the complex and strain-specific MaxOD patterns observed across strains.

A

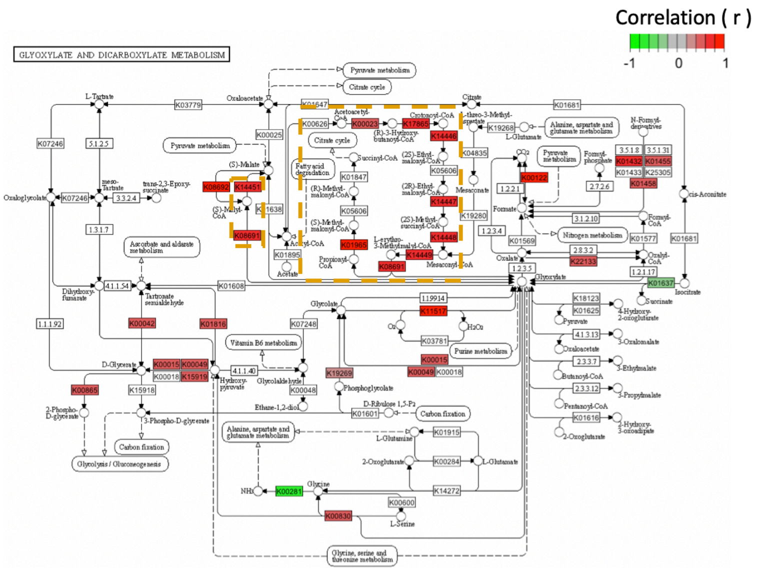


B

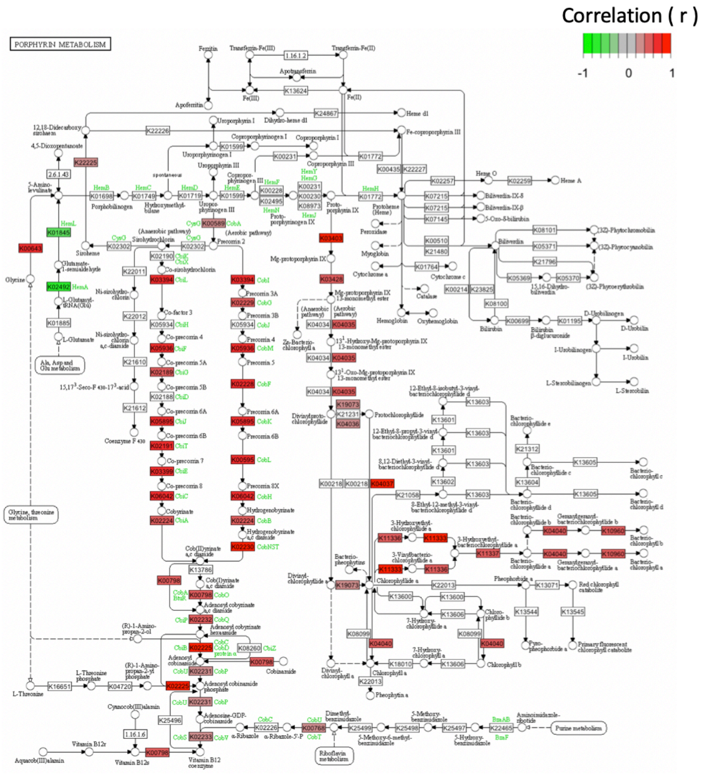


**Figure S5**. KEGG pathway enrichment for growth on organic acids. Genes highly correlated with growth on organic acids are mapped onto the Glyoxylate and Dicarboxylate **(A)** and the Porphyrin and Chlorophyll **(B)** Metabolism KEGG pathways; the colors represent the sign and strength of the correlation, ranging from green (-1) to red (1). In **(A)**, the portions of the KEGG pathway representing the Ethylmalonyl-CoA pathway are denoted by the dashed gold boxes.

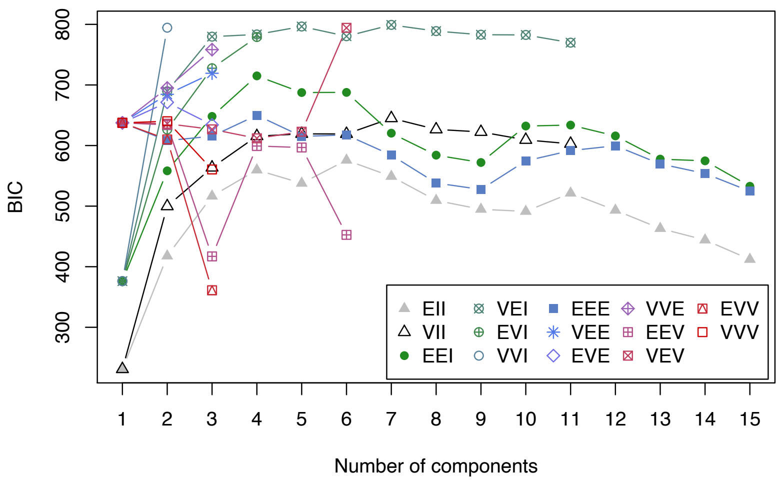


**Figure S6.** Gaussian Mixture Model (GMM) model selection. The Bayesian information criterion (BIC) is plotted as a function of the number of clusters for 14 different parameterization approaches implemented by mclust (shown in subset, see (68) for details). The model with the highest BIC is identified as having the optimal parameters; in our case, VEV (Volume, Shape, Orientation = Variable, Equal, Variable) with seven clusters.
